# The genomic signature and transcriptional response of metal tolerance in brown trout inhabiting metal-polluted rivers

**DOI:** 10.1101/2024.05.30.595956

**Authors:** Josephine R Paris, R Andrew King, Joan Ferrer Obiol, Sophie Shaw, Anke Lange, Vincent Bourret, Patrick B Hamilton, Darren Rowe, Lauren V Laing, Audrey Farbos, Karen A Moore, Mauricio A Urbina, Ronny van Aerle, Julian M Catchen, Rod W Wilson, Nicolas R Bury, Eduarda M Santos, Jamie R Stevens

## Abstract

Industrial pollution is a major driver of ecosystem degradation, but it can also act as a driver of contemporary evolution. As a result of intense mining activity during the Industrial Revolution, several rivers across the southwest of England are polluted with high concentrations of metals. Despite the documented negative impacts of ongoing metal pollution, brown trout (*Salmo trutta* L.) survive and thrive in many of these metal-impacted rivers. We used population genomics, transcriptomics, and metal burdens to investigate the genomic and transcriptomic signatures of potential metal tolerance. RADseq analysis of six populations (originating from three metal-impacted and three control rivers) revealed strong genetic substructuring between impacted and control populations. We identified selection signatures at 122 loci, including genes related to metal homeostasis and oxidative stress. Trout sampled from metal-impacted rivers exhibited significantly higher tissue concentrations of cadmium, copper, nickel, and zinc, which remained elevated after 11 days in metal-free water. After depuration, we used RNAseq to quantify gene expression differences between metal-impacted and control trout, identifying 2,042 differentially expressed genes (DEGs) in the gill, and 311 DEGs in the liver. Transcriptomic signatures in the gill were enriched for genes involved in ion transport processes, metal homeostasis, oxidative stress, hypoxia and response to xenobiotics. Our findings reveal shared genomic and transcriptomic pathways involved in detoxification, oxidative stress responses, and ion regulation. Overall, our results demonstrate the diverse effects of metal pollution in shaping both neutral and adaptive genetic variation, whilst also highlighting the potential role of constitutive gene expression in promoting metal tolerance.

## Introduction

Pollution causes long-term environmental impacts with severe implications for biodiversity and human health (Schwarzenbach et al., 2010; Travis & Hester, 1991), yet growing evidence of tolerance to toxic chemicals has resulted in a greater awareness of pollution in mediating contemporary evolution (Hendry et al., 2017; Smith & Bernatchez, 2008; Whitehead, 2014). Whilst organisms can respond to short-term stress via phenotypic plasticity, understanding the mechanisms underpinning tolerance to chronic exposure requires consideration of long-term adaptive responses (Badyaev, 2005; Ghalambor et al., 2007). However, because pollutants are often not acutely toxic by design, have wide-ranging physiological effects, and occur in complex mixtures in the environment (involving different metals in varying concentrations and forms, which can interact with each other), dissecting the role of pollutants in mediating adaptive responses can be difficult, especially in natural populations (Whitehead et al., 2017).

Metals are naturally occurring elements, but are found at significantly higher concentrations in ecosystems associated with certain human activities such as mining, agriculture and industrial processes (Vareda et al., 2019). These activities have led to elevated levels in aquatic environments, in some cases exceeding the mortality thresholds for many species (see Kumar et al., 2019). Essential metals, such as copper, iron, manganese and zinc, are crucial for life, acting as key components for various biological processes including development, energy metabolism and immunity, but can become toxic when threshold concentrations are exceeded (Chandrapalan & Kwong, 2021). On the other hand, non-essential metals, such as arsenic and cadmium, lack known biological function, and thus toxicity manifests when such metals are present in the environment in particular chemical states at lower concentrations (Wood et al., 2012a). Metals exert their toxic effects through different mechanisms, including the production of reactive oxygen species (ROS), DNA damage, and the disruption or inhibition of enzymes, proteins, and subcellular organelles (Hartwig et al., 2002; Valko et al., 2005; Vangronsveld & Clijsters, 1994). Therefore, adaptation to metal polluted environments would require physiological mechanisms that reduce toxicity while maintaining levels of essential metals.

Over the last four decades, a wealth of information has been published on the impact of metals on fish including: discerning the role of different tissues in metal uptake, storage and detoxification (*e.g.* Bradley & Morris, 1986; Eastwood & Couture, 2002); understanding the physiological processes regulating homeostasis and toxicity (Shahjahan et al., 2022; Wood et al., 2012a, 2012b); and characterising the molecular mechanisms underlying metal homeostasis (Hamilton & Mehrle, 1986; Srikanth et al., 2013). At the genomic level, studies have evaluated the impact of metal pollution on genetic structure and diversity (Bourret et al., 2008; Maes et al., 2005), as well as genetic adaptation (Hamilton et al., 2017; Laporte et al., 2016). Further, a number of studies have demonstrated that metal pollution also affects epigenetic processes, with long-term impacts on the health of individuals and populations (Abdelnour et al., 2024; Pierron et al., 2023). Meanwhile, at the transcriptomic level, several studies have quantified differences in gene transcription between metal-impacted and control fish (Bélanger-Deschênes et al., 2013; Coffin et al., 2022; Pierron et al., 2009; Uren Webster et al., 2013), highlighting genes and pathways potentially involved in the mitigation of metal toxicity. However, measuring gene expression from individuals sampled directly from their native sites does not allow for a distinction between short-term responses to acute stress, versus responses to chronic stress (López-Maury et al., 2008). In this regard, constitutive gene expression, *i.e.*, the higher baseline expression of certain genes or pathways, can confer protection to stressors by providing a pre-emptive adaptive response (Hendry, 2016; Rivera et al., 2021). Investigations aimed at understanding the impacts of metal pollution, and the evolution of potentially adaptive responses, would benefit from a multifaceted approach combining physiology, genomics and transcriptomics.

Brown trout (*Salmo trutta* L.) in the southwest of England provide one of the best purported cases of metal tolerance in aquatic environments. Mining in the area dates back to the Bronze Age (ca.L2500-800 BCE), and later, during the Industrial Revolution (1850-1900), the region dominated the global output of tin, copper and arsenic (Burt et al., 2014; Timberlake, 2017). As a testament to its role in shaping Britain’s industrial development and its broader influence on the global mining industry, the area has been designated as a UNESCO World Heritage Site (The Cornwall and West Devon Mining Landscape; https://www.cornishmining.org.uk/). The longevity and intensity of mining has resulted in elevated contemporary metal concentrations in several rivers in the region (Rainbow, 2020), and consequently, negative ecological and biological effects including a reduction or total extermination of aquatic life (Brown, 1977; Bryan et al., 1987; Hart et al., 2020). Yet brown trout are abundant, and are able to reproduce and survive in many of these rivers, despite the fact that they experience both chronic and seasonally-varying acute exposure to metals at concentrations known to be lethal to naïve salmonids (see Table S1). Based on neutral genetic markers, trout from metal-impacted rivers were shown to be genetically differentiated from trout sampled from nearby control rivers, and displayed significantly reduced genetic diversity (Osmond et al., 2024; Paris et al., 2015). Another study showed that trout from metal-impacted rivers had significantly elevated tissue-metal burdens (Uren Webster et al., 2013). This opens the question of how these brown trout cope with metal toxicity. Transcriptomics analysis comparing metal-impacted trout and control fish sampled directly from the river highlighted differentially expressed genes involved in metal and ion homeostasis, oxidative stress, and immunity (Uren Webster et al., 2013). However, the extent to which these signatures represent a short-term response to metal toxicity is unknown. To investigate the signatures of potentially adaptive responses, genetic variation should be surveyed using genome-wide markers, and transcriptional responses should be quantified in fish outside of the metal-impacted environment.

Here, we investigate the signatures of potential long-term adaptation to metal pollution in wild brown trout using a combination of genomics, transcriptomics, and tissue-metal burdens. First, we use RADseq to genotype six brown trout populations (originating from three metal-contaminated and three control rivers) to assess how metal pollution has shaped the neutral and non-neutral genetic make-up of metal-impacted populations. We hypothesised that if metals have caused population-level impacts, we would observe distinctive patterns of genetic variation in metal-impacted populations. We also reasoned that if metal pollution has driven an adaptive response, selection signatures at outlier loci related to metal handling will be evident. Second, we aimed to evidence and assess, if and how metals are absorbed into the tissues of metal-impacted brown trout. We address this by quantifying differences in metal burdens across five tissues (gill, liver, gut, kidney, and muscle) between metal-impacted trout and those from a matched control site. To refine our understanding of metal handling, we quantified tissue-burdens from individuals sampled directly from the river, and after an 11-day depuration in control (metal-free) water. Finally, we quantified gene expression differences between metal-impacted and control trout following the 11-day period of depuration in two key tissues known for their roles in metal uptake and detoxification (gill and liver) to assess the molecular processes involved in potential long-term responses to metal pollution. Integrating across these approaches provides a broad understanding of adaptation to metals, and of the mechanisms underpinning evolution to environmental stress in wild fish.

## Materials and Methods

### 2.1 Study Sites and Sampling

The Cornwall and West Devon (United Kingdom) mining region contains many small rivers affected by metal pollution. While many rivers are affected by high concentrations of metals, individual rivers are polluted with different mixtures of metal pollutants (Figure 1A). For clarity, we term trout sampled from these rivers as “metal-impacted”. To reduce confounding effects of biogeographic history, we sampled what we term “control” trout from rivers with lower concentrations of metals sampled within the same region. Due to the nature of the bedrock and extensive history of mining in the region, we note that control rivers may also have been historically impacted by metals, but in recent years show comparatively lower metal concentrations (Table S2).

**Figure 1.**
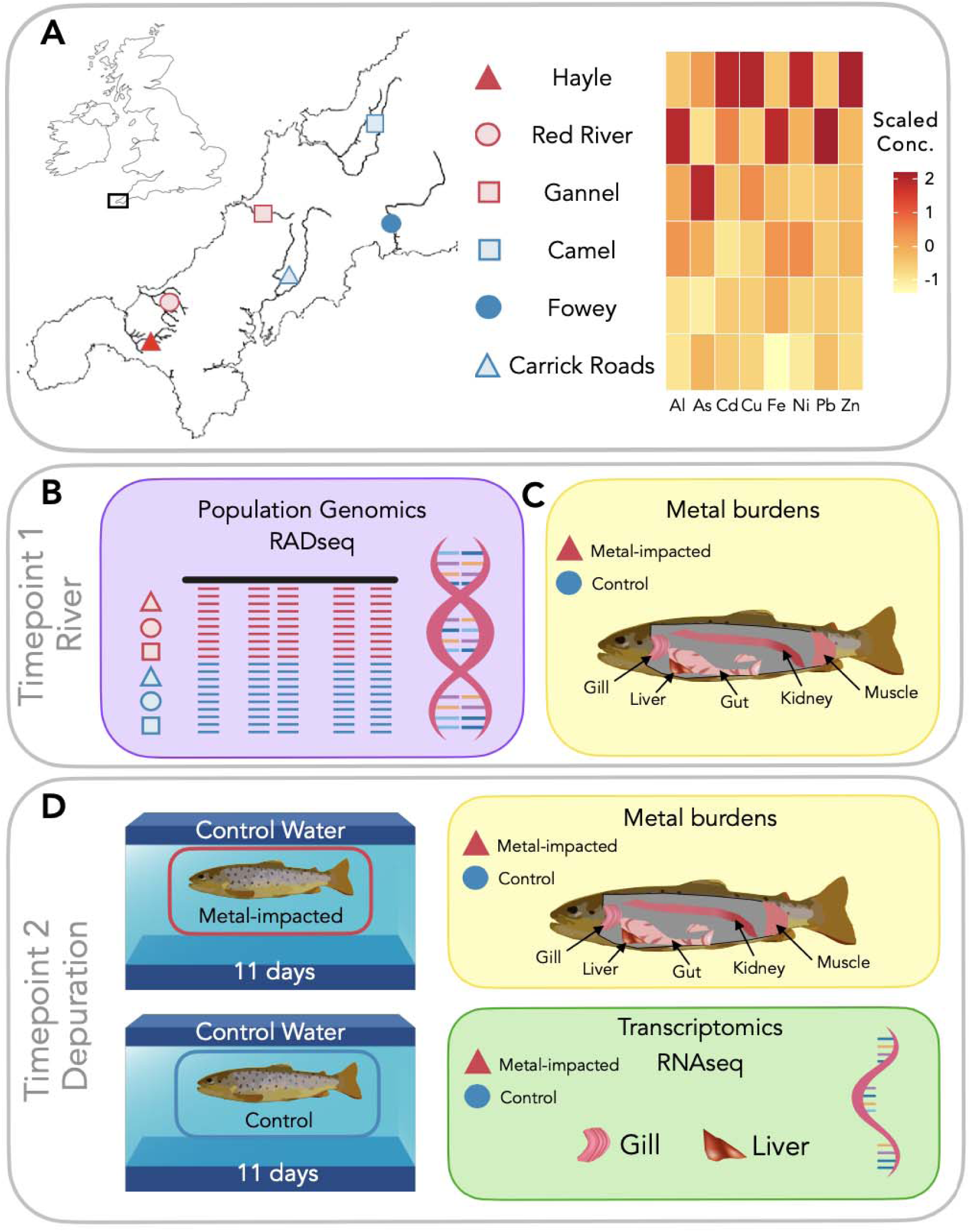
Schematic of collection sites and experimental design. (**A**) Location within the UK of the Cornwall and West Devon mining region and location of sampling sites. Metal-impacted sites are coloured in red and control sites are coloured in blue. Filled coloured shapes represent sites sampled for tissue-metal burden analysis before and after a period of depuration. The heatmap shows the scaled metal concentrations of each river (averaged over a 15-year period; full data available in Table S2) (**B**) Timepoint 1: River. For population genomics, *n* = 20 brown trout were sampled from three metal contaminated rivers: Hayle (red triangle); Red River (red circle); Gannel (red square) and three control rivers: Camel (blue square); Fowey (blue circle) and Carrick Roads (blue triangle). (**C**) Timepoint 1: River. For tissue burden analysis, metal-impacted trout were sampled from the Hayle (red triangle; *n* = 10) and control trout were sampled from the Fowey (blue circle; *n* = 10). (**D**) Timepoint 2: Depuration. Trout were sampled after 11 days of depuration in control water for tissue burden analysis (*n* = 10 per group) and transcriptomic analysis of the gill and liver using RNAseq (*n* = 6 per group).

For the population genomics analyses, a total of *n* = 120 trout were sampled across 14 sampling sites from a total of six rivers (*n* = 20 per river; Figure 1A, B; Table S2; Table S3). The largest adult fish (> 150 mm fork-length) were sampled to ensure resident origin (*i.e.*, not anadromous individuals straying from nearby rivers) and prolonged survival in the metal-polluted rivers. Fish were anaesthetised using a stock solution of benzocaine (100 g/L in ethanol) diluted to 50 mg/L in river water prior to adipose fin-clip removal. Tissue samples were stored in 95% ethanol at 4° C prior to DNA extraction.

For the quantification of tissue burdens and transcriptomics analysis, trout (0+ parr) were sampled from the River Hayle (metal-impacted, *n* = 26) and the River Fowey (control, *n* = 26). Fish from the River Hayle were chosen as representative metal-impacted trout as they showed the most distinct genetic signatures in our analyses, suggesting minimal gene flow with populations from neighbouring rivers over multiple generations. This population has also been characterised and studied previously (Minghetti et al., 2014; Osmond et al., 2024; Paris et al., 2015; Uren Webster et al., 2013). Trout from the River Fowey were selected as control fish since the river exhibits relatively low metal contamination and the most comparable water chemistry (which is a key determinant of metal bioavailability) to that of the River Hayle (Table S4).

All fish were caught by backpack electrofishing, following Home Office (UK) guidelines, Environment Agency (UK) authorisation, and relevant landowner permissions.

Trout were transported to the Aquatic Resources Centre, University of Exeter, in water from their river of origin. Immediately upon arrival (on the same day of sampling), we randomly selected 10 metal-impacted and 10 control fish (from the n = 26 fish sampled per site) to represent the status of individuals in the river (Timepoint 1: River; Figure 1C). After 11 days of depuration and acclimation to laboratory conditions (in control water), we randomly sampled 10 metal-impacted and 10 control fish (Timepoint 2: Depuration; Figure 1D). The 11-day depuration period was chosen based on a balance between metal elimination kinetics (*i.e.*, rate constants; see Veltman et al., 2008) and the challenges of maintaining wild brown trout in aquaria.

Fish were humanely killed via an overdose of benzocaine (0.5 g/L), followed by destruction of the brain. We recorded the wet body mass and fork length, and the condition factor was calculated (*K*) = [mass (g) x 100] / [fork length (cm)^3^] before dissecting and weighing tissues for analysis. As a growing number of studies have demonstrated sex-related differences in ecotoxicological responses (e.g. Burger et al., 2007), we attempted to account for any potential sex-related biases in our metal-burden and RNAseq analyses. Due to the young age of the fish, gonadal sex could not be reliably assessed, so additional muscle tissue was collected and stored in 95 % ethanol at −20 °C for genetic sex determination (King & Stevens, 2020). All animal work was carried out in accordance with the UK Animals Scientific Procedures Act (ASPA) 1986 and was ethically approved by the Animal Welfare and Ethical Review Body at the University of Exeter.

### 2.2 Laboratory depuration and acclimation: water chemistry and fish maintenance

In the laboratory, 7-9 fish were housed in 39 L glass tanks. For the depuration period, fish were held in synthetic control (metal-free) water to mimic the water chemistry of the River Hayle (Minghetti et al., 2014) using salts of the following concentrations: 391 µM NaCl, 128 µM KHCO_3_, 500 µM MgSO_4_, 350 µM Ca(SO_4_), 75 µM Ca(NO_3_)_2_. Salts were made up to a 50x stock concentration, dosed into constant-flowing reverse osmosis purified water using a conductivity pump set to dose between 265 and 270 μS/cm. Water chemistry of the synthetic freshwater was assessed via daily pH and conductivity measurements. Freshwater was delivered via a flow-through system at a flow rate of 100 ml/min. The laboratory was temperature controlled at 15°C and a natural day-night 12h:12h light cycle was used. Individuals were fed daily to satiation.

### 2.3 Population genomics using RADseq

#### 2.3.1 DNA extraction and RADseq library preparation

DNA was extracted from adipose fin tissue using the QIAGEN DNeasy Blood & Tissue Kit (QIAGEN). DNA concentrations were determined using the QuantiFluor ONE dsDNA assay system (Promega). Single-digest *SbfI* RADseq libraries were prepared following the protocol described by Etter et al. (2011). Libraries were amplified using 12 PCR cycles and were sequenced on an Illumina HiSeq 2500 (100bp single-end) at a target coverage of 20x per individual.

#### 2.3.2 RADseq data pre-processing

Data were processed using Stacks v2.6 (Rochette et al., 2019). Raw reads were demultiplexed and cleaned using *process_radtags*. Optimised parameters for construction of *de novo* RAD loci were determined using the *r80* method (Paris et al., 2017). Seven of the 120 individuals were removed due to low coverage. We assembled loci *de novo* (minimum stack depth (m) = 3, distance between stacks (M) = 2, distance between catalog loci (n) = 2) and then aligned loci to the brown trout genome (Ensembl: version fSalTru1.1; Hansen et al., 2021) using *stacks-integrate-alignments* (Paris et al., 2017) to provide positional information. We used stringent parameters for filtering the loci, in order to detect top candidates for potential metal adaptation. Loci were filtered with the following parameters in *populations*: *-r 80, -p 6, -min-maf 0.1, -max-obs-het 0.8*. For analyses sensitive to linkage, we used the *– write-single-snp* option and pruned loci in linkage (LD dataset) using Plink v1.9 (Purcell et al., 2007).

#### 2.3.3 Population structure, genetic diversity, and effective population size estimation

Population structure was assessed by Principal Components Analysis (PCA) and the neighbour-joining (NJ) tree method. PCA was conducted in Plink v1.9 (Purcell et al., 2007). For constructing the NJ tree, the VCF file was converted to Phylip format using *vcf2phylip* (Ortiz, 2019), requiring a minimum of 100 samples per SNP. Distances were calculated using the *dist.dna* function in ape v5.6 (Paradis & Schliep, 2019), using the K80 substitution model and removing pairwise missing data.

Expected heterozygosity (H_E_), observed heterozygosity (H_O_) and the inbreeding coefficient (*F*_IS_) were calculated for each population using Hierfstat (Goudet, 2005). To assess variance in individual genetic diversity, we calculated the proportion of polymorphic sites across positions that were variable in at least one population using the --het option in VCFtools v0.1.16 (Danecek et al., 2011). Pairwise Weir and Cockerham’s *F*_ST_ were computed in Hierfstat (Goudet, 2005). To perform an assessment of fixed alleles within each population, we calculated allele frequencies using VCFtools v0.1.16 (Danecek et al., 2011) and plotted the number of fixed alleles per population in R v4.0 (R Core Team, 2021).

Biases in estimating the effective population size (*N*_e_) using linkage-disequilibrium methods are well-known (see Waples, 2024), especially when using RADseq and low sample sizes (Marandel et al., 2020). To alleviate these biases, we used the recently described methods implemented in currentNE, which performs particularly well in the small sample size scenario (Santiago et al., 2024). currentNE was run by estimating the average number of full siblings (k parameter) from the input data (a Hardy-Weinberg equilibrium and missing data filtered VCF file per population), using 40 chromosomes as estimated from the brown trout reference genome. 90% confidence intervals were provided by the neural network (ANN).

#### 2.3.4 Identification of loci associated with metal pollution and water chemistry

To identify candidate genes potentially involved in metal adaptation, we used a combination of statistical methods to detect loci showing high allele frequency differentiation: pcadapt v4.3.3 (Luu et al., 2017) and OutFLANK v0.2 (Whitlock & Lotterhos, 2015), and genotype-environment association (GEA) tests: Redundancy Analysis (RDA; Capblancq et al., 2018) and Latent Factor Mixed Models (LFMM v1.5; Frichot et al., 2013). pcadapt was run using *K* = 4, as identified using the *screeplot* function. p-values were corrected for multiple testing using an alpha value of 0.05. For OutFLANK, left and right trim fractions were set to 0.35 and 0.06, respectively. We removed low heterozygosity variants (< 0.1) and used a q-value threshold of 0.05.

For GEA tests, we used 41 water chemistry and metal contamination variables for each of the six rivers (14 sample sites; Table S3). We used the average value for each variable (data covers a 15-year period; Table S2). Due to microgeographic differences in metal contamination within each river, we extracted allele frequency data for each of the 14 sampling sites to match the environmental data (4664 loci).

RDA was run adapting the approaches described by Bourret et al., (2014) and Forester et al., (2018). To avoid collinearity amongst the environmental predictors, the environmental data were reduced to 10 Rotated Components (RC) by running a PCA (10 factors, *varimax* rotation). RCs were reduced to a number of factors identified by a parallel analysis criterion (Horn, 1965) using 1000 bootstraps. We used a 0.01 significance threshold (0.68 loading cutoff) to associate an environmental parameter to a factor. Following the RDA, a global analysis of variance (*anova.cca*; 999 permutations) was performed to assess the global significance of the RDA and an analysis by axis was run to determine if retained RCs were significantly correlated with population-level allele frequencies. Candidates were identified as variants in the tails of the RDA constrained axis distribution. A cutoff of 2.5 standard deviations (two-tailed p-value of 0.012) was used.

LFMM was run using the RCs generated above. The number of latent factors was estimated using the *snmf* function (*K* = 4). We ran 10,000 iterations with a burn-in of 5000. The Gibbs sampler was run five independent times on the dataset in order to estimate the appropriate average LFMM parameters (z-score values). p-values were adjusted using estimation of the genomic inflation factor (λ) for each RC variable. Significance was tested using an FDR < 0.05.

Candidate loci (*i.e.*, all loci identified from the analyses described above) were assessed for functional gene-level effects using the Variant Effect Predictor tool in Ensembl (McLaren et al., 2016). We also classified the potential gene-level effects of the fixed alleles in each of the metal-impacted populations. Variants were classified as intergenic, intronic, synonymous, and non-synonymous, including the potential variant consequences.

### 2.4 Tissue-metal burden

#### 2.4.1 Laboratory analysis

Five tissues (gill, liver, gut, posterior kidney, and muscle) were dissected, freeze-dried and weighed, and stored at −20 °C. Tissues were digested in 500μl of nitric acid (70 %, purified by redistillation, ≥ 99.999 % trace metals basis, Sigma Aldrich) for 48 hours at room temperature (∼20 °C), followed by digestion with 0.1 % hydrogen peroxide (Fisher; Hydrogen Peroxide 30% (w/v) Extra Pure SLR) for a further 24 hours. Digested tissue was diluted 1:10 in ultrapure deionised H_2_0 and acidified with 0.1 % nitric acid. A standard of 10 μg/L yttrium (Y) was added to each sample as an internal standard to perform within-sample corrections. Metal concentrations (quantified as ng/mg) were measured by Inductively Coupled Plasma Mass Spectrometry (ICP-MS) (E:AN 6100DRC, Perkin-Elmer, Cambridge, UK).

#### 2.4.1 Statistical analysis

Data were checked for normality (Shapiro test) and homogeneity of variance (Levene test) using the stats package (v3.6.1) in R v4.0 prior to conducting statistical tests and were log transformed when necessary. Assessment of burdens for each tissue were statistically assessed using a two-way ANOVA, with ‘metal-impacted’ and ‘control’ fish specified as Group, and Timepoint (‘River’, or ‘Depuration’) specified as Factor. For analyses where statistical differences were identified, post-hoc Tukey tests were performed to conduct pairwise multiple comparisons, accepting a significance threshold of FDR ≤ 0.05.

### 2.5 Transcriptomics analysis

#### 2.4.1 RNA extraction, library construction, and sequencing

Gill and liver tissues were dissected from euthanised fish (*n* = 6 metal-impacted; *n* = 6 control) and were snap-frozen in liquid nitrogen and stored at −80°C prior to analysis. To avoid sex-biassed gene expression differences, we selected three (genetically-determined) females and three males from each group. RNA was extracted using a QIAGEN RNeasy extraction kit (QIAGEN), including an on-column DNAse digestion step. RNA quantification and quality assessments (accepting a RIN > 8.0) were performed using a Bioanalyser (Agilent RNA 6000 Nano). Libraries were prepared using the Illumina TruSeq Stranded mRNA library preparation protocol. Concentrations of the cDNA libraries were quantified using a Tapestation (D1000 ScreenTape) and final library pool concentrations were determined by qPCR. Libraries were sequenced (125 bp paired-end) on an Illumina HiSeq 2500.

#### 2.4.2 RNAseq data pre-processing

Raw reads were cleaned using Trimmomatic v0.27 (Bolger et al., 2014) and aligned to the brown trout reference genome (Ensembl version fSalTru1.1; Hansen et al., 2021) using STAR v2.5.4 (Dobin et al., 2012). Transcript abundances were calculated at gene regions using featureCounts v2.0.1 (Liao et al., 2014). Genes with low read counts were removed by only including genes with more than 10 counts in ≥ 6 samples. This reduced the initial dataset of 43,946 genes to 28,261 and 21,820 expressed genes in the gill and liver, respectively.

#### 2.4.2 Differential gene expression analysis

Analysis of differentially expressed genes (DEGs) was performed using DESeq2 v1.40.2 (Love et al., 2014). For assessing clustering of the biological replicates, we performed PCA on a variance stabilising transformed (VST) normalised count matrix. Size factors and dispersion were estimated using the default DESeq2 parameters. A negative binomial GLM was used to test for DEGs, retaining only those genes with a significance threshold FDR ≤ 0.05 and log2foldchange values of ≤ −1 or ≥ 1.

All genes (including non-significant DEGs, but with dispersion outliers removed) were used to test for a statistical enrichment in Gene Ontology (GO) terms via Gene Set Enrichment Analysis (GSEA) using clusterProfiler v3.16.1 (Yu et al., 2012). Lists containing GO categorisation of brown trout terms (using the Brown Trout Genes fSalTru1.1 database; AH85410) were downloaded from Ensembl using BioMart v2.44 (Smedley et al., 2009).

Genes were ranked by a single ranking metric by multiplying the log fold-change by the negative logarithm (base 10) of the adjusted p-value. This was done to ensure that genes with both large fold-changes and low p-values receive higher combined ranks. Minimum and maximum gene set sizes of 20 and 200 were used. GO terms with a Benjamini–Hochberg adjusted p-value ≤ 0.25 were classified as being significantly enriched or underrepresented.

## Results

### 3.1 Population genomics

#### 3.1.1 RADseq libraries and loci construction

We generated an average of 2,279,882 reads (SEM ± 119,860) per individual at an average of 20X coverage, assembling 10,115 SNPs in 8,718 loci genotyped across 80% of the individuals from each of the six rivers. Details of the RADseq libraries, including number of reads and coverage obtained for each individual are shown in Table S5.

#### 3.1.2 Population genetics characterisation of metal-impacted and control trout

Overall, we observed strong genetic sub-structuring with the highest pairwise *F*_ST_ values detected in comparisons to metal-impacted trout populations, in particular those sampled from the River Hayle (*F*_ST_ > 0.1 in all comparisons) and the lowest pairwise *F*_ST_ values (range: River Camel-River Fowey *F*_ST_ = 0.015 to Carrick Roads-River Fowey; *F*_ST_ = 0.03) were observed between control trout populations (Figure 2A).

**Figure 2.**
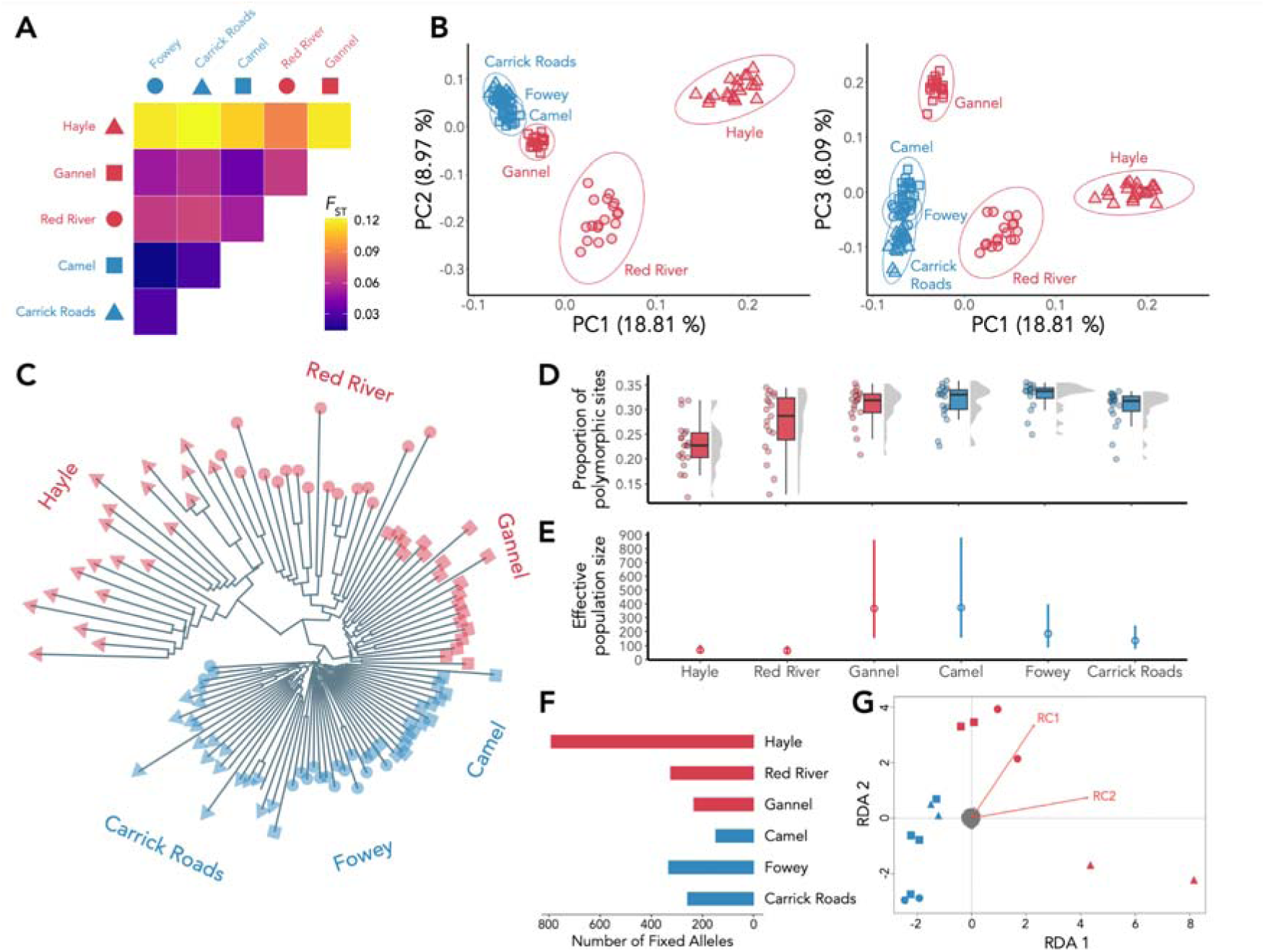
Population genomic data describing the structure and diversity of metal-impacted and control trout. (**A**) Heatmap of pairwise *F*_ST_ calculated between all sampling sites. (**B**) Principal Component Analysis (PCA) showing PC1 (∼19% of the variance) against PC2 (∼9% of the variance) and PC1 against PC3 (∼8% of the variance). Individuals are coded by sampling site as in panel A. (**C**) Neighbour-joining tree from aligned SNPs, computed using the K80 substitution model. (**D**) Proportion of polymorphic sites for each individual sampled from each of the six sampling sites. (**E**) Estimates of the effective population size (*N*_e_) for each of the six sampling sites. (**F**) Quantification of fixed alleles calculated at each of the six sampling sites. (**G**) Redundancy Analysis (RDA; *p* = 0.007, R^2^ = 0.2145) showing RDA1 (*p* = 0.037) and RDA2 (*p* = 0.1). Red vectors represent RC1 (water chemistry features; Table S7) and RC2 (Cd, Cu, Ni, Zn). In all panels, sampling sites are coloured by metal-impacted (red) or control (blue). For panels A, B, C, and F the sampling sites are coloured and shaped as: Hayle (red triangle); Red River (red circle); Gannel (red square); Camel (blue square); Fowey (blue circle) and Carrick Roads (blue triangle).

PCA separated the metal-impacted trout populations from one another, and also from the control trout (Figure 2B). PC1 (18.8%) defined trout from the River Hayle, PC2 (8.97%) separated the Red River and River Gannel trout. PC3 (8.09%) further separated the River Gannel trout and also revealed patterns of IBD in the control trout populations. In the NJ tree, metal-impacted trout showed three distinctive branches, with the River Hayle and Red River individuals originating from a shared branch in the NJ tree, whereas the River Gannel formed its own group (Figure 2C). The River Hayle and Red River showed particularly long branches, indicative of higher substitution rates. We also observed genetic substructuring within the River Hayle (upstream and downstream of a highly metal-contaminated middle region; Durrant et al., 2011; Paris et al., 2015) and the Red River (by tributary) in the NJ tree.

We observed differences in the genetic diversity and effective population size (N_e_) estimates between metal-impacted and control trout (Figure 2D; Figure 2E). Compared to control sites, genetic diversity was significantly lower (Mann-Whitney U *p* < 0.005) in metal-impacted trout sampled from the Red River (H_O_ = 0.28) and the River Hayle (H_O_ = 0.23), but not in the River Gannel (H_O_ = 0.31). The proportion of polymorphic sites also showed a more even distribution in the control trout, whereas long tails were observed in the River Hayle and Red River, indicative of high variance among the individuals sampled from these sites (Figure 2D). Further investigation of the heterozygosity per sampling site showed that this high variance was not caused by microgeographic differences in the upstream and downstream reaches of the River Hayle, or by the two sampled tributaries in the Red River. Differences in observed heterozygosity were reflected in the inbreeding coefficient (*F*_IS_), which was overall higher in metal-impacted trout compared to control trout, being highest in trout sampled from the River Hayle (*F*_IS_ = 0.25), and lowest in the River Fowey (*F*_IS_ = 0.04).

*N*_e_ estimates (Figure 2E) for the River Hayle (*N*_e_ = 68; 90% CI: 43-106) and Red River (*N*_e_ = 64; 90% CI: 40-100) were low, but the River Gannel (*N*_e_ = 366; 90% CI: 157-863) had a higher *N*_e_ than the River Fowey (*N*_e_ = 186; 90% CI: 87-398) and Carrick Roads (*N*_e_ = 137; 90% CI: 77-243), and was comparable to that of the River Camel (*N*_e_ = 372; 90% CI: 157-881). Overall, metal-impacted trout did not show a higher number of fixed alleles, except for the River Hayle, which showed a much higher number compared to trout from all other sites (Figure 2F).

#### 3.1.3 Identification of adaptive loci

Using four methods, we identified a total of 122 candidate loci potentially involved in metal adaptation (Table S6). Of the pcadapt candidates (*n* = 40), the majority were associated with PC1 (*n* = 18, which separates the River Hayle). Remaining outliers were associated with PC2 (*n* = 14; separates the Red River), and PC3 (*n* = 8; separates the River Gannel). OutFLANK identified eight candidates, which were primarily driven by divergent *F*_ST_ values observed in comparisons to the River Hayle.

The RDA was globally significant (*p* = 0.007; R^2^ = 0.2145) and both axes were also significant (RDA axis 1: *p* = 0.037; RDA axis 2: *p* = 0.1, accepting a significance threshold ≤ 0.1; Figure 2G). We identified a total of 30 loci which were significantly associated with these axes (two-tailed *p* = 0.012). RDA axis 1 (five loci) separated the River Hayle, and to a lesser extent, the Red River and River Gannel from the control group. RDA axis 2 (25 loci) separated the Red River and River Gannel populations from the control group, suggesting this axis represents loci which are diverging in these populations, but not in the River Hayle. Correlations between the loadings on the RCs and the RDA axes showed that RDA axis 1 was primarily associated with the environmental features of RC2: Cd, Cu, Ni and Zn (Table S7), which are the primary metal pollutants of the River Hayle. RDA axis 2 was primarily associated with the environmental features of RC1 (water chemistry features; Table S7). Using LFMM, we uncovered significant associations of 44 loci (RC1: 22 loci; RC2: 14 loci; RC3: 8 loci. We found no overlap amongst the loci identified using the four outlier approaches.

Of the 122 candidates, 60 loci showed good-quality alignments (MQ=60) to genes in the brown trout genome (Table S6). Twenty-eight of the variants were intronic, 21 variants were located upstream or downstream of genes, three were synonymous variants, and nine variants had functional gene-level predictor effects. Gene functions were varied, but we identified candidates potentially involved in metal homeostasis, and response to oxidative stress and xenobiotics, as well as genes involved in calcium homeostasis, tumour regulation, transcriptional regulation, and immunity (see Discussion). Of the 1348 fixed variants identified in each of the metal-impacted populations, 42 aligned to genes in the brown trout genome. All variants were found to be intronic, except for a missense variant in the River Hayle, located in an open reading frame (Table S8). Amongst the intronic variants, we identified a gene involved in protein ubiquitination (*usp2b*), a potassium channel regulator (*kcnab1b*), an iron ion binding gene (*iscub*), and an aryl hydrocarbon receptor protein (BMAL2).

### 3.2 Laboratory maintenance and fish morphometrics

In the laboratory, all fish were observed feeding; there were no mortalities and no changes in behaviour were observed. Stable water chemistry of the synthetic freshwater was maintained throughout the experiment (Table S9). Metal-impacted trout were significantly larger than control fish; however, there was no significant difference in condition factor (Table S10). The metal-impacted and control groups showed uneven numbers of males and females and this was taken into consideration during the statistical comparisons (Timepoint 1: metal-tolerant M:F 2.67, control M:F 1.5; Timepoint 2: metal-tolerant M:F 1.4, control M:F 1.17).

### 3.3 Tissue-metal burden analysis

No differences were observed between Timepoint 1 and Timepoint 2 for any metal in any tissue in the control trout. Sex did not affect the statistical significance of the metal-burden data. Full statistical results for the metal-burden data can be found in Table S11.

At Timepoint 1 (river), gill metal concentrations were 7-, 6-, 1- and 2-fold higher in metal-impacted trout compared to control trout for Cd (*p* < 0.0001), Cu (*p* < 0.0001), Ni (*p* = 0.01) and Zn (*p* < 0.0001), respectively (Figure 3A). In metal-tolerant trout, the concentrations of Cd, Cu and Ni decreased during the depuration period (Timepoint 2) (Cd, *p* = 0.0003; Cu, *p* < 0.0001; Ni, *p* = 0.01), but Zn did not (*p* = 1). Although the concentration of Cd decreased in Timepoint 2, it was still higher than the concentrations found in control trout at the same timepoint (*p* < 0.0001). Zn burdens also remained higher than control trout at Timepoint 2 (*p* < 0.0001).

**Figure 3.**
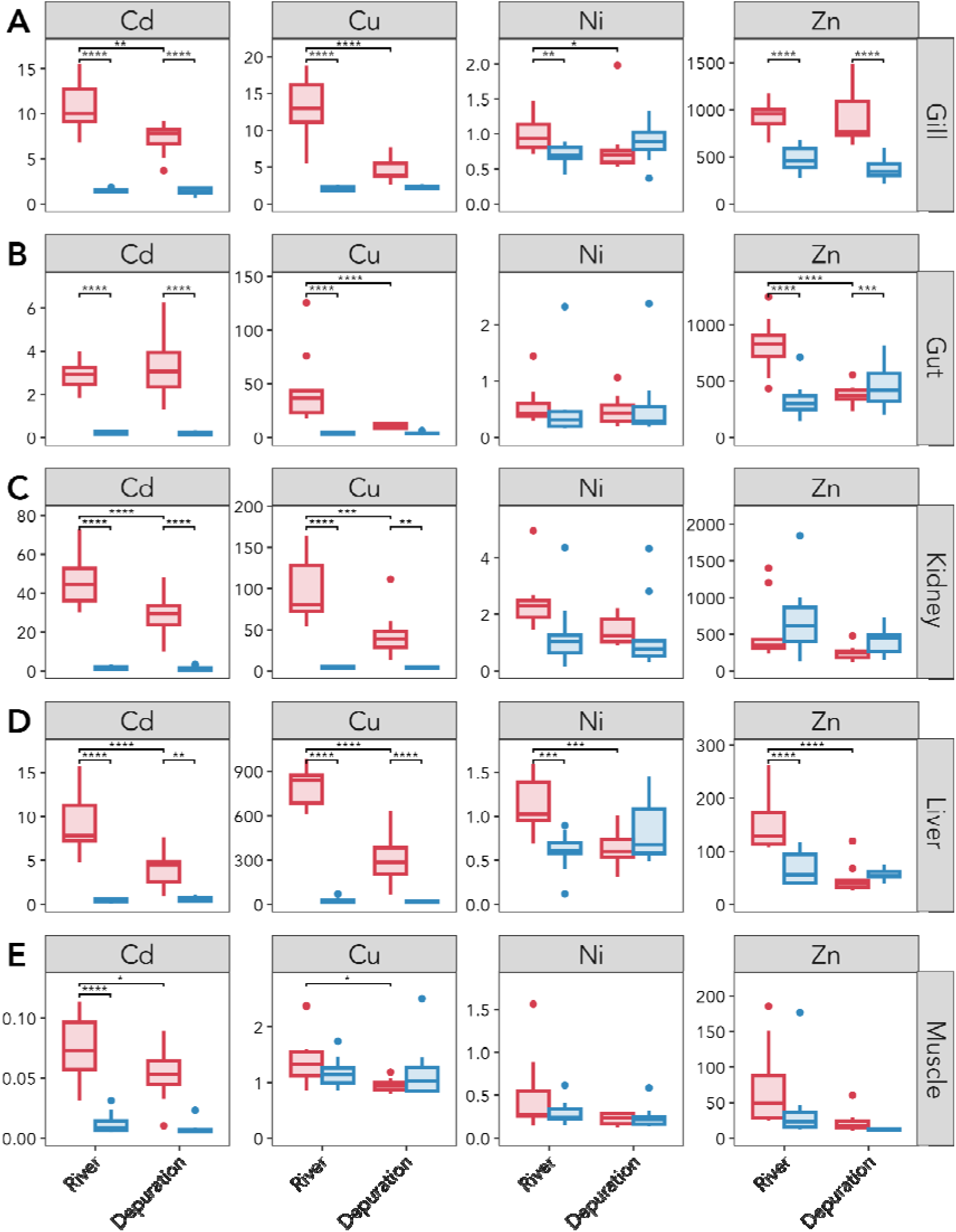
Tissue-metal burden data of cadmium (Cd), copper (Cu), nickel (Ni), and zinc (Zn) measured in metal-impacted (in red) and control (in blue) trout measured at Timepoint 1 (River) and Timepoint 2 (Depuration). Metal burden is measured as ng/mg of dry weight tissue for the (**A**) Gill; (**B**) Gut; (**C**) Kidney; (**D**) Liver; and (**E**) Muscle. Significance was measured using a Two-way ANOVA (full details in Table S11). Only significant results from pairwise Tukey tests are included where **** indicates p≤0.0001, *** indicates p≤0.001, ** indicates p≤0.01, and * indicates p≤0.05.

In the gut, metal burdens at Timepoint 1 were 14-, 11- and 2-fold higher in metal-impacted trout for Cd (*p* < 0.0001), Cu (*p* < 0.0001) and Zn (*p* < 0.0001), respectively (Figure 3B). Ni burden was not higher (*p* = 0.99) and did not decrease at Timepoint 2 (*p* = 0.98). The concentrations of both Cu and Zn decreased at Timepoint 2 (Cu, *p* < 0.0001, Zn, *p* < 0.0001) to concentrations comparable to those observed in control trout (Cu, *p* = 0.83; Zn, *p* = 0.68). Cd concentrations did not decrease (*p* = 0.56), with 18-fold higher concentrations being measured in metal impacted fish compared to control trout after Timepoint 2 (*p* < 0.0001).

In the kidney, metal burdens measured in fish at Timepoint 1 were 27- and 23-fold higher in metal-impacted trout for Cd (*p* < 0.0001) and Cu (*p* < 0.0001) (Figure 3C). Although both metals showed a significant depuration (Cd, *p* < 0.0001; Cu, *p* = 0.0002), they remained 17- and 11-fold higher than control trout at Timepoint 2 (Cd, *p* < 0.0001; Cu, *p* = 0.006). Ni was higher in metal-impacted trout, but this was marginally non-significant in comparison to control trout (*p* = 0.09). No statistical differences were observed for Zn.

In the liver, metal burdens were 21-, 30-, 2-, and 2-fold higher in metal-impacted trout at Timepoint 1 for Cd (*p* < 0.0001), Cu (*p* < 0.0001), Ni (*p* = 0.001) and Zn (*p* < 0.0001), respectively (Figure 3D). Metal-impacted trout showed a depuration of all metals at Timepoint 2 (Cd, *p* < 0.0001, Cu, *p* < 0.0001, Ni, *p* = 0.002; Zn, *p* < 0.0001). However, concentrations of Cd and Cu remained 7- and 17-fold higher than control trout at Timepoint 2 (Cd, *p* = 0.003; Cu, *p* < 0.0001).

In the muscle, metal-impacted trout showed 6-fold higher Cd burden concentrations compared to control trout (Cd, *p* < 0.0001) when measured at Timepoint 1, and Cd did not depurate (Cd, *p* = 0.05) (Figure 3E). Although slightly higher burdens of Cu were observed in metal-impacted trout, this was not significant (*p* = 0.41); Cu did however depurate (*p* = 0.03). Ni and Zn did not show any significant differences.

### 3.3 Comparison of the transcriptional landscapes between metal-impacted and control trout following a period of depuration

#### 3.3.1 RNAseq libraries and gene expression profiles

Across the six samples per group (metal-impacted and control), we generated an average of 16,549,620 (SEM ± 1,341,005) reads for the gill and 11,588,849 (SEM ± 1,329,081) reads for the liver (Table S12). PCA revealed a clear separation between metal-impacted and control trout in their transcription profiles after depuration. In the gill, replicates from each group were separated along PCA axis 1 (49% of the variation; Figure 4A), whereas in the liver they were separated along PCA axis 2 (21% of the variation; Figure 4B).

**Figure 4.**
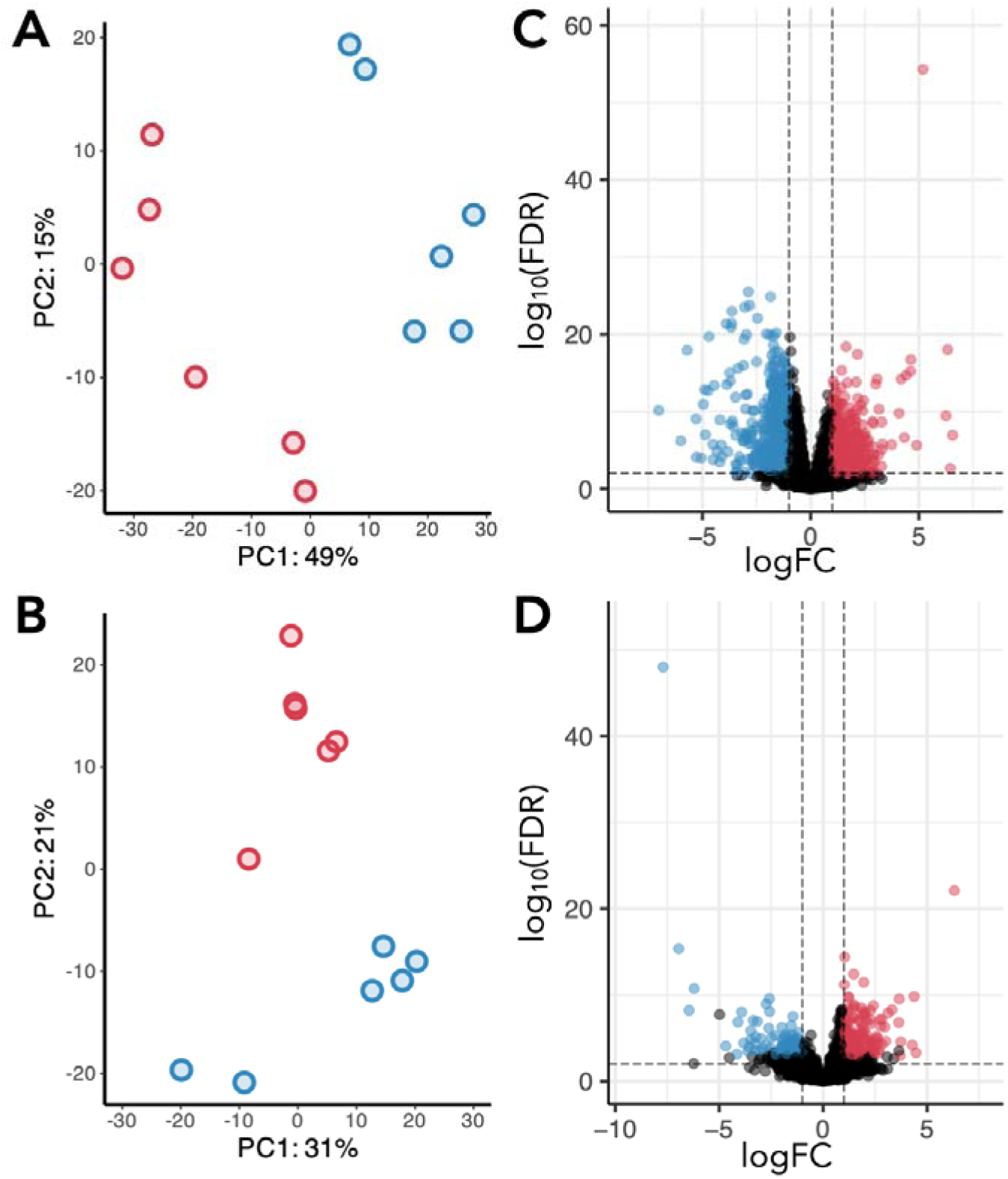
Differential gene expression between metal-impacted and control trout. (**A**) Principal Components Analysis (PCA) of the variance stabilising transformation (VST) normalised counts in the gill. (**B**) Principal Components Analysis (PCA) of the VST normalised counts in the liver. Coloured dots represent the transcriptomic profile of individual fish from the metal impacted (red) and control (blue) rivers. (**C**) Volcano plot of the 28,261 genes measured in the gill. (**D**) Volcano plot of the 21,820 genes measured in the liver. Coloured dots in the volcano plots correspond to differentially expressed genes (DGEs) with FDR ≤ 0.05 and a log2foldchange of 1 (red) or −1 (blue), meaning genes are over-expressed, or under-expressed in metal-impacted trout, respectively.

#### 3.3.2 Identification of differentially expressed genes

Analysis of the differentially expressed genes (DEGs) revealed 2,042 DEGs (FDR ≤ 0.05) in the gill, of which 1,180 were over-expressed (log2FC ≥ 1) and 862 were under-expressed (log2FC ≤ −1) in metal-impacted trout compared to control trout (Figure 4C; see Table S13 for the full list of significant DEGs). We identified several metal-binding and antioxidant genes that remained over-expressed after depuration in the gill tissue of metal-impacted trout, including metallothionein 2 (*mt2*), metal response element binding transcription factor 2 (*mtf2*), metalloreductase (*steap4*), methionine sulfoxide reductase B2 (*msrb2*), thioredoxin-related transmembrane protein 2 (*tmx2a*), three glutathione-dependent enzymes (*tstd1*, *mgst3*, *gstz1*), and lysyl oxidase (*loxl2a*, *loxl2b*). We also identified genes with well-known roles in response to oxidative stress and response to xenobiotics, including an epoxide hydrolase, two cytochrome p450 genes, five cytochrome c oxidase subunits, a cytochrome b-245 chaperone (*cybc1*), both aryl hydrocarbon receptors (*ahr1a*, *ahr1b*), and fourteen heat-shock proteins. Related specifically to the known activity of the gill in response to metals, we also identified ATP-binding cassette (ABC) transporters (*abca4a*, *abcg2a*, *abcc5*), and two transcripts encoding carbonic anhydrase 4 (*ca4*). Amongst the under-expressed genes, we observed a high number of immune-system related genes, including interferon-induced proteins, interleukins, and E3 ubiquitin protein ligases.

In the liver, 311 DEGs (FDR ≤ 0.05) were identified, of which 200 were over-expressed (log2FC ≥ 1), and 111 were under-expressed (log2FC ≤ −1) in metal-impacted trout compared to control trout after the depuration period (Figure 4D; see Table S14 for the full list of DEGs). We did not observe the same level of transcriptional changes in the liver, with very few over-expressed metal-binding, oxidative stress response, or xenobiotic handling genes. However, the metal-binding protein *heavy metal-binding protein HIP-like*, the oxidative stress gene superoxide dismutase 3 (*sod3*), and the well-known xenobiotic metabolism gene, cytochrome P450 1A (*cyp1a*) were over-expressed in the metal-impacted group.

Intersecting the gill and liver DEGs, we identified 25 shared over-expressed genes, and eight shared under-expressed genes (Table S15). Of note, *DENN domain-containing protein 5B-like* was over-expressed in both tissues and was the most strongly over-expressed gene in both tissues.

#### 3.3.3 Gene Set Enrichment Analysis (GSEA) on Gene Ontology (GO) terms

To further investigate the transcriptomic landscapes of metal-impacted trout following depuration, we explored enriched and underrepresented GO terms via GSEA. In the gill, we identified several enriched biological processes indicative of changes in ionoregulation including chloride, calcium, sodium, and potassium ion transporters, and response to oxidative stress, including regulation of small GTPase mediated signal transduction.

Underrepresented biological processes included microtubule movement, DNA repair and replication, immune responses, protein folding, catabolism, and ubiquitination. Amongst the enriched molecular function pathways, we identified ion channel activity, including voltage-gated potassium and calcium channel activity, metallocarboxypeptidase activity, cytochrome-c oxidase activity, and copper ion binding and iron sulphur cluster binding. The identified GO terms associated with both enriched and underrepresented processes in the gill dataset are detailed in Table S16.

In the liver, we found a weak concerted biological signal related to metal tolerance. We found no significantly enriched or underrepresented biological pathways, but cytochrome-c oxidase activity was a significantly enriched molecular function. The identified GO terms associated with both enriched and underrepresented processes in the liver dataset are detailed in Table S17.

## Discussion

We applied an integrative approach, using genomics, transcriptomics, and tissue-metal burdens, to investigate the impacts of, and potential long-term adaptation to metal pollutants in wild brown trout. Population genomics analysis confirmed that metal-impacted trout are highly genetically differentiated from control trout, and that two of the three populations show significantly lower genetic diversity and low *N*_e_, highlighting the effects of metal pollutants on the genetic make-up of these populations. Using a series of outlier tests, we identified selection signatures for loci associated with metal homeostasis, and response to oxidative stress and xenobiotics. Measured immediately after removal from the river, metal-impacted trout showed considerably higher concentrations of metals across multiple tissues, suggesting the requirement of mechanisms to cope with increased metal burden. After 11 days in control water, patterns of metal depuration were tissue- and metal-specific, pointing towards a complex network of biological responses involved in metal handling. Analysis of gill transcriptional profiles revealed over-expressed genes and enriched biological processes in metal-tolerant trout involved in ionoregulation, metal homeostasis and oxidative stress providing a mechanistic understanding of metal tolerance.

### 4.1 Evidence of high tissue concentrations of metals in metal-impacted trout

Tissue-metal burden analysis in individuals sampled at their river of origin showed significantly higher concentrations of metals in the tissues of metal-impacted trout compared to control fish, similarly to that reported in Uren Webster et al., (2013) on fish collected from comparable locations. The varying levels of metal concentrations across the different tissues points to the diverse roles of these organs in organismal level metal uptake, storage, and detoxification. The gills, being in intimate contact with the water, and the gut, via ingestion of metal-contaminated prey, are well-characterised as the main routes of metal uptake in fish (Wood, 2012). In the gills, we found elevated concentrations of all metals measured in this study (Cd, Cu, Ni and Zn), pointing to this tissue as the main entry site. Meanwhile, elevated concentrations of Cd, Cu and Zn in the gut suggests brown trout are also acquiring metals from ingested food in the river. However, Ni was not elevated in the gut; previous research has shown that pre-exposure to Ni reduces gastrointestinal uptake, suggesting a homeostatic interaction might exist for this metal (Chowdhury et al., 2008).

The liver and kidney provide key functions in the detoxification and excretion of metals in fish (Wood, 2012). All metals measured (Cd, Cu, Ni and Zn) were significantly elevated in the liver, supporting its role as the main tissue of metal storage and detoxification in vertebrates. In contrast, only Cd and Cu were elevated in the kidney. As reported here, the kidney has been previously found to accumulate the highest concentrations of Cd (Andres et al., 2000; De Smet et al., 2001). Muscle generally shows a weak accumulating potential for metals (Jia et al., 2017), and we found that only Cd was significantly elevated within this tissue.

Overall, the observation of elevated tissue burdens of metals in trout living in metal-impacted environments supports the hypothesis that physiological mechanisms, likely influenced by genomic and/or transcriptomic mechanisms must be involved in the dynamics of internal metal storage and detoxification, and in preventing the harmful effects associated with such high metal loads.

### 4.2 Metal pollutants impact the genetic structure and reduce genetic diversity of metal-impacted trout

We observed strong genetic sub-structuring between metal-impacted and control trout across all analyses. Such patterns have been observed previously using both neutral microsatellites (Paris et al., 2015) and SNPs (Osmond et al., 2024). Despite their geographic proximity, metal-impacted trout populations showed strong genetic separation, both from one another, and also from control trout. In contrast, differentiation was low among control trout occupying clean rivers, which are geographically much more distant from one another. For example, the mouths of the River Hayle and Red River (metal-impacted) are separated by just ∼9 km, whereas the River Camel and the Carrick Roads (control) are separated by ∼270 km of coastline. Population genetic studies on salmonids often report a high incidence of population structure (Bekkevold et al., 2020; King et al., 2020; King et al., 2024), frequently attributed to processes of local adaptation (Fraser et al., 2011; Primmer, 2011; Taylor, 1991). However, identifying what proportion of the genetic structure is explained solely by local adaptation is confounded by neutral demographic processes. Yet it can also be argued that if populations are geographically close, but experience substantially different environmental pressures, the expected differentiation due to non-selective factors is small (Wadgymar & DeMarche, 2022; Whitlock & Lotterhos, 2015). The low genetic differentiation observed among control populations also supports the local adaptation hypothesis. Moreover, there are no known physical barriers separating the mouths of the metal-impacted rivers, and thus the strong population structure could also indicate that non-adapted individuals which stray into metal-impacted rivers suffer adverse toxic effects (Sergeant et al., 2022). Over time, a reduction in gene flow would be expected to further increase genetic drift and strengthen genetic differentiation.

The effect (neutral and/or non-neutral) of metal pollutants was particularly marked in the River Hayle, where we observed micrographic structuring based on metal contamination levels, with trout sampled upstream and downstream of an extremely toxic middle section of the river (∼1.5km), showing strong genetic differentiation (Paris et al., 2015). As well as indicating that metal toxicity may act as a chemical barrier to migration within the river (Hecht et al., 2007; Williams & Gallagher, 2013), these microgeographic differences could also indicate finer-scale adaptive responses. Indeed, primary gill cell cultures showed higher metallothionein gene expression patterns when exposed to River Hayle water downstream from the toxic section (Minghetti et al., 2014).

Metal-impacted populations do not appear to be experiencing the same genetic impact in response to metal pollutants. The River Hayle, the Red River, and the River Gannel, are all impacted by different combinations of metal pollutants and also differ in their water chemistry features (which affect metal bioavailability). This result could stem from a number of factors: metal toxicity at different concentration thresholds within each river; disparate adaptive responses in each of the metal-impacted populations; differences in the population size and demographic history of these populations. Previous analysis using Approximate Bayesian Computation (ABC) showed that bottlenecks occurred in all metal-impacted populations (Paris et al., 2015). The River Hayle and Red River trout appeared to be most affected by metal pollution, with these populations showing the strongest population structure and particularly long branches in the neighbour-joining tree. The latter observation indicates higher substitution rates in these populations, which can be the result of multifarious biological processes, including higher levels of inbreeding, drift, episodes of heightened adaptive evolution, but also higher mutation rates, or differences in life-history traits such as generation time (Mendes & Hahn, 2016). Increased mutation rates have been evidenced in metal-impacted sunfish (*Lepomis auratus*; Theodorakis et al., 2006), and metal-impacted yellow perch (*Perca flavescens*) had lower longevity than control conspecifics (Couture & Pyle, 2008). Genetic diversity and *N*_e_ were also lower in the River Hayle and Red River populations, but particularly so in the former, which also had a higher inbreeding coefficient. Such observations support the hypothesis that environmental pollution can cause significant genetic erosion (van Straalen & Timmermans, 2002). Nevertheless, the fact that trout remain in these rivers, and show evidence of extremely high concentrations of metals in their tissues, suggests an adaptive component to metal tolerance.

### 4.3 Assessment of candidate loci involved in local adaptation to metal pollution

To detect candidate loci underpinning potential metal tolerance we applied several statistical tests, each with different assumptions, using a particularly rigorously set of filtered loci and stringent cut-off criteria for significance. This was to reduce spurious correlations between loci, environmental variables, and the strong population structure we observed (Ahrens et al., 2018; Hoban et al., 2016). Amongst the 122 outlier loci, we identified 12 variants with potential gene-level effects, several of which have functional annotations with potential relevance to metal adaptation. We identified a splice-gene variant in *map35k*, which is known to be activated by various stressors including ROS and calcium overload (Ichijo et al., 1997) and is regulated in response to copper (O’Doherty et al., 2020). In addition, we identified a synonymous variant in *scara3* which is involved in scavenging oxidative molecules (Han et al., 1998; Sone et al., 2010) and a missense variant in *ferritin, heavy subunit-like* which is involved in the storage of iron in a nontoxic state (Munro & Linder, 1978), both potentially contributing to a reduction of oxidative stress caused by elevated intracellular metal ion concentrations. We also identified a stop-loss mutation in *cyp7a1*, one of the cytochrome P450 enzymes involved in mono-oxygenation reactions in a wide range of endogenous and exogenous compounds (Uno et al., 2012); and which has been shown to be regulated in response to toxic metals (Chi et al., 2019). Genes involved in transcriptional regulation were also identified including a 3’-UTR variant in ZNF239 and a missense variant in PCGF3. Further, we identified a splice donor variant in *hmmr*, which regulates cell growth (He et al., 2020) and is modified in exposure to arsenic (Dodmane et al., 2013), and missense variants in *mfhas1* and *inavaa* which both function in innate immunity: *mfhas1* is a regulator of *tlr2* and *tlr4* (Shi et al., 2017; Zhong et al., 2015), and *inavaa* is stimulated by pattern recognition receptors including *tlr2* and *tlr4* (Yan et al., 2017). Whilst these genes represent promising candidates underpinning local adaptation to metals, we recognise that we have not surveyed the full variation potentially responsible for metal tolerance due to, for example, limitations in the RADseq approach (Lowry et al., 2017, but see Catchen et al., 2017), and neglecting the role of structural variants (SVs; Mérot et al., 2020), especially duplications arising from the salmonid whole-genome duplication event (Moriyama & Koshiba-Takeuchi, 2018). Further research using whole-genome sequencing methods are required to increase our understanding of potential candidates involved in adaptation to metals in these trout populations.

### 4.4 Metal- and tissue-specific depuration patterns over time

We observed both metal- and tissue-specific patterns of depuration, pointing to differential mechanisms for each of the measured metals for deposition and excretion. The patterns of depuration in the gill, liver and kidney are of particular interest. In the gill, Cd, Cu and Ni decreased in concentration over time, but Zn did not, and although Cd depurated from the gill, its concentration remained higher than that observed in control fish. Toxic concentrations of all metals can have deleterious effects on the gill, including oxidative stress (Sevcikova et al., 2011), impaired ionoregulation (Wood et al., 2012a, 2012b), damage and mucus accumulation (Playle, 1998), necessitating their rapid detoxification and removal (Saibu et al., 2018). Rapid clearance (within 16h) of Cd and Cu from the gill has been reported previously in rainbow trout (*Oncorhynchus mykiss*), with accompanying increases in the liver (Handy, 1992). Indeed, metal-impacted trout continued to show high burdens of all metals in the liver, but especially Cd and Cu, both of which showed evidence of depuration, but not to comparable concentrations observed in the control fish. We observed similar patterns for Cd and Cu in the kidney, suggesting overall that the gill is essential in detoxifying these metals, and that they are subsequently transported to the liver and kidney for excretion (Garai et al., 2021). We also observed a high retention of Cd across all tissues, which has been observed previously, and over much longer time periods than our experiment allowed for (Defo et al., 2015; Kraemer et al., 2005).

As others have shown, these results indicate that tissue metal concentrations can be high, but the metals within the tissue are likely present in a non-toxic form (*e.g.*, Campbell et al., 2008; Leonard et al., 2014; Shekh et al., 2021; Wang & Rainbow, 2006). The physiological processes governing this are attributed to subcellular partitioning, whereby metals are compartmentalised into either a metal-sensitive fraction (MSF), which is considered the target of attack by metals in cells, or the biologically detoxified metal (BDM) portion, which is considered to alleviate toxicity (see review by Wang, 2013). Notably, there is evidence that over time, metals are moved from the MSF to the BDM, and that fish surviving long-term in chronically metal-impacted environments have partitioned the majority of their tissue metal burdens into the BDM (Wood et al., 2012b). Overall, the observation of tissue-specific shifts in metal burdens, as well as the retention of high metal burdens among particular tissues, emphasises the importance of detoxification and excretion in metal-impacted brown trout, necessitating molecular mechanisms underpinning these processes.

Our study design used only two timepoints to protect the welfare of experimental fish (Díaz et al., 2020), particularly as we were removing wild fish from relatively small natural populations. This strategy limited our ability to quantify depuration over both: i) additional timepoints, which would have allowed a more detailed insight into the journey of metal depuration; and ii) a longer depuration period, permitting a more confident determination of “true” depuration (including our insights into constitutive gene expression). However, we highlight the challenges of maintaining wild fish under laboratory conditions (Huntingford et al., 2006), including acclimation to captivity (Baillon et al., 2016), and perhaps, more importantly, uncertainty regarding whether metal-impacted trout would ever completely depurate to reach the background tissue-metal burdens observed in the control population.

### 4.5 Constitutive gene expression as a potential mechanism of adaptation to metal pollution

We observed distinct transcriptional profiles between metal-impacted and control trout after 11 days of depuration, suggesting a role for constitutive gene regulation in mitigating metal toxicity. In particular, the gill had a higher number of DEGs (2,042), 58% of which were over-expressed in comparison to the control trout, including those involved in metal detoxification, oxidative stress and response to xenobiotics. These included the well-known metal-binding gene metallothionein (*mt2*), a key protein involved in the homeostasis and detoxification of Cd, Cu and Zn (Hamilton & Mehrle, 1986). In a previous study quantifying transcriptional differences measured from trout sampled directly from the river, *mt2* was over-expressed in both the gill (log2FC: 8.2) and liver (log2FC: 5.6) (Uren Webster et al., 2013), whereas after depuration, we found that this gene was over-expressed (albeit to a lesser extent) only in the gill (log2FC: 1.6). We also identified genes with well-known roles in stress and xenobiotic response, including heat-shock proteins (Gupta et al., 2010). We further identified the overexpression of both isoforms of the aryl hydrocarbon receptor (*ahr1a*, *ahr1b*), which mediate the biochemical response to toxic compounds including metals (reviewed in Kou et al., 2024) and have been implicated in toxicant adaptation in killifish (*Fundulus heteroclitus*; Aluru et al., 2015; Whitehead et al., 2017) and Atlantic tomcod (*Microgadus tomcod*; Wirgin et al., 2011). In our RADseq data, we also identified an intronic SNP in an aryl hydrocarbon receptor protein. Related specifically to the activity of the gill in response to metals, we identified the overexpression of several ABC transporters, which are known to be involved in the uptake, transfer and elimination of xenobiotics (Kropf et al., 2020). We observed little overlap in the DEGs identified in the gill and liver, pointing to the distinctive roles of these tissues in metal homeostasis. However, the most highly over-expressed gene in both tissues was *DENN domain-containing protein 5B-like*. The DENN domain is an evolutionarily conserved module found in all eukaryotes (Zhang et al., 2012). DENN domain-containing proteins directly interact with Rab GTPases (Marat et al., 2011), acting as molecular switches in the regulation of multiple intracellular trafficking processes (Homma et al., 2021), that respond to stress (Ganotra et al., 2023) and interact with metallothionein (Knipp et al., 2005; Wolff et al., 2008).

Overall, despite the high number of identified DEGs, we identified a relatively low number of significantly enriched or underrepresented GO terms. This could indicate low concordance in the function of the genes which govern the recovery process from metal exposure. Although we sampled individuals of the same age, and of equal sex ratios (three males and three females per group), this result could also indicate the stochasticity in using wild fish in our experiment, and moreover, that individual wild trout might not experience metal toxicity in the same way, nor bear the same metal tolerance phenotype. Finally, variability in the process of acclimation to laboratory conditions could also confound the identification of metal detoxification pathways. Related to this, although we identified some GO terms related specifically to metal homeostasis in the gill, we identified fewer than anticipated, and the liver showed very little signal related to metal homeostasis. A recent study in mosquitofish (*Gambusia affinis*) inhabiting metal-polluted environments, also found that many DEGs were not associated with known metal response pathways, highlighting the multifariousness of metal response pathways and indicating that undocumented pathways could be playing a role in metal tolerance (Coffin et al., 2022). Generally, these results highlight the complexity of understanding metal tolerance, and calls attention to our lack of knowledge surrounding adaptation and acclimation to metals in wild vertebrates.

In the gills, we identified a number of pathways involved in ion transport. An important mechanism of metal toxicity is the disruption of ion transport pathways in the gill, which eventually causes disruptions of ionic balance in the body (Evans, 1987; Wood, 1992). In particular, Cd and Zn are known to share uptake routes in the gill (Ca transport pathways), where acute toxicity occurs as a result of the disruption of Ca homeostasis (Hogstrand et al., 1996; Niyogi & Wood, 2004). On the other hand, Cu uptake in the gill is known to occur mainly via Na transport pathways, and acute toxicity manifests via the disruption of Na homeostasis (Grosell & Wood, 2002). We identified Ca ion transmembrane transport, voltage-gated calcium channel activity, and Na ion transport as significantly enriched processes in the gill. Additionally, metals can cause the inhibition of the Na+/K+-ATPase pump (Glover et al., 2016; Grosell & Wood, 2002; Haverinen & Vornanen, 2023), and Cu can increase Cl loss (Mazon et al., 2002). Both potassium and chloride ion transporters were enriched, as well as voltage-gated potassium channels. We also identified an intronic variant in a potassium channel regulator (*kcnab1b*) in the RADseq data. Together, this suggests that compensatory increases in ion transport activity may underpin responses to limit the toxic effects of metals in the gill of metal-impacted trout.

Our results suggest that constitutive gene expression is likely an important adaptive mechanism permitting metal tolerance. Plasticity theory predicts that when populations experience consistent inducing conditions over multiple generations, a plastic response can eventually become constitutively expressed, even in the absence of the original cues inducing the change (*i.e.* genetic assimilation; Pigliucci et al., 2006; Waddington, 1961). It has also been suggested that constitutive expression can provide higher fitness than short-term responsive expression (Evans, 2015; Geisel, 2011; Snell-Rood et al., 2010). We note that we are limited in fully deciphering what proportion of the DEGs are constitutively expressed solely due to metal handling, as the DEG signatures could also be indicative of response to other environmental factors. However, the clear separation of the gene expression profiles, the identification of relevant genes and pathways in the gill, accompanied by the observed differences in tissue-metal concentrations, and their depuration, support an important role of constitutive gene expression in promoting metal tolerance.

## 5 Considerations and conclusions

We provide new insights into the evolution of metal tolerance in brown trout originating from historically metal-polluted rivers. The observation of elevated tissue-metal burdens confirms that metal pollution affects the trout inhabiting these rivers and supports the hypothesis that adaptive strategies to cope with the insult of elevated metals within internal tissues have evolved. Across our genomic and transcriptomic analyses, we identified signatures of genetic differentiation in metal-impacted rivers and a number of expressed or positively selected genes, which are functionally linked to known metal response molecular pathways. Specifically, several RADseq outlier loci were identified as genes with known roles in the response to oxidative stress and we also identified transcriptomic pathways associated with this process, including the regulation of small GTPases (Ferro et al., 2012), heat shock proteins (Kalmar & Greensmith, 2009), AhR pathway activation (Kou et al., 2024), and cytochrome-c oxidase activity (Srinivasan & Avadhani, 2012).

The high number of differentially expressed genes in the gill, accompanied by the strong patterns of metal accumulation and depuration, highlights this organ as being of key functional importance for metal tolerance in this population with other organs playing also a role in the dynamics of metal homeostasis. Overall, our data contributes to an in-depth understanding of the molecular pathways allowing brown trout to live in highly contaminated environments and the evolutionary processes driving these adaptations.

## Data Availability Statement

De-multiplexed RADseq data for all samples are available in the European Nucleotide Archive (ENA) under the project accession: PRJEB78481. Note that data for the following samples (cam01, cam02, fal01, gan01, gan02, hay06 - hay13) were published in Paris et al. 2017 (doi.org/10.1111/2041-210X.12775) and are also available at the NCBI under BioProject PRJNA379215. RNAseq data is available from ArrayExpress accession E-MTAB-10209.

## Author contributions

JRP, RWW, NRB, EMS and JRS conceived the study, designed, and performed research. JRP, RAK, SS, AL, VB, JFO, PBH, DR, LVL, MAU, RvA and JMC analysed data. KAM and AF contributed analytical tools. JRP, EMS and JRS wrote the paper with contributions from co-authors. All authors read the manuscript and provided comments.

## Acknowledgements and Funding

The authors dedicate this article to Professor Mike Bruford, who was the external viva voce examiner of the PhD thesis of JRP, and who offered valuable advice on the research. Environmental data were provided by the Environment Agency Devon and Cornwall Teams. Fieldwork, electrofishing, permits and access to sites was planned and undertaken with assistance from the Westcountry Rivers Trust, in particular Matthew Healey and Giles Rickard. Fieldwork was assisted by Janice Shears, Philip Shears and Adam Porter. The UK Environment Agency provided assistance in the collection of water chemistry features and selection of appropriate sampling sites. Simon Toms (UK Environment Agency) granted permission to collect wild brown trout for the physiology experiment. Aquatic Research Centre (ARC) staff at the University of Exeter provided assistance with fish husbandry and experimental design, in particular Gregory Paull and Charles Tyler who oversaw the ethical aspects of the laboratory experiment and who provided guidelines on Home Office Regulations. Jennifer Fitzgerald, Rebecca Millard and Hannah Littler assisted with experimental sampling. RAD-seq and RNA-seq sequencing was performed at the University of Exeter Sequencing Service, which utilised equipment funded by the Wellcome Trust Institutional Strategic Support Fund (WT097835MF), Wellcome Trust Multi-user Equipment Award (WT101650MA) and BBSRC LOLA award (BB/K003240/1). Research was co-funded by the Environment Agency, the Westcountry Rivers Trust, and the University of Exeter.

